# rSWeeP: A R/Bioconductor package deal with SWeeP sequences representation

**DOI:** 10.1101/2020.09.09.290247

**Authors:** Danrley Fernandes, Mariane G. Kulik, Diogo J. S. Machado, Jeroniza N. Marchaukoski, Fabio O. Pedrosa, Camilla R. De Pierri, Roberto T. Raittz

## Abstract

The rSWeeP package is an R implementation of the SWeeP model, designed to handle Big Data. rSweeP meets to the growing demand for efficient methods of heuristic representation in the field of Bioinformatics, on platforms accessible to the entire scientific community. We explored the implementation of rSWeeP using a dataset containing 31,386 viral proteomes, performing phylogenetic and principal component analysis. As a case study we analyze the viral strains closest to the SARS-CoV, responsible for the current pandemic of COVID-19, confirming that rSWeeP can accurately classify organisms taxonomically. rSWeeP package is freely available at https://bioconductor.org/packages/release/bioc/html/rSWeeP.html.

## 1 Introduction

In the era of Big Data, vector and alignment-free approaches to compare and represent biological sequences stand out for being more efficient than most alignment-based heuristic methods^[1]^. Studies show that vector representation of biological data presents an effective solution for the Bioinformatics field^[2–5]^.

Recently, SWeeP model presented expressive results in the analysis of complete proteomes. SWeeP is a model implemented in MatLab, which allows the representation of biological information through the projection of DNA sequences in vectors with reduced dimensionality^[5]^. Here, we present rSWeeP, an implementation of SWeeP model using R/Bioconductor programming language. To test the effectiveness of the implementation, we run a performance test, phylogenetic analysis and Principal Components Analysis (PCA) using viral proteomes as a dataset. Our results showed that rSWeeP maintains the effectiveness of SWeeP model in comparison of large numbers of sequences. In the case study, we identified the proximity between the virus responsible for the current SARS (Severe acute respiratory syndrome) pandemic (COVID19) and the virus responsible for the SARS pandemic in 2003, suggesting that it is the same coronavirus, as already reported in other study^[6]^.

## 2 Methods

All steps of rSWeeP implementation were performed on an Intel core I5 320Gz with 12 GB of RAM.

### Implementation

rSWeeP input is a multiFASTA file or an “AAStringSet” class object, containing amino acid sequences. rSWeeP consists in two main functions:

1. “OrthBase”: to generate an orthonormal matrix in a specified size;
2. “SWeeP”: to generate and project SWeeP vectors to compare the sequences information.

### Dataset

We used all viruses’ sequences in the “complete genome” category on the NCBI assembly, available at https://ftp.ncbi.nlm.nih.gov/genomes/Viruses/ (downloaded on February 6th, 2020). A tutorial for running rSWeeP and the trees are available at https://github.com/DanrleyRF/Suplementar. For all analysis, we applied the same protocol adopted in the study by De Pierri and colleagues (2020) ^[5]^.

## 3 Results and Discussion

### Construction of Phylogenetic Trees

To explore the potential of the rSWeeP package, we created SWeeP vectors from the protein annotations of all the complete viral genomes available at NCBI, in total, 31,386 proteomes sequences (31k). A viral phylogenetic tree was built almost ten times larger than the reference found in the literature^[3]^.

There are many specimens for each virus species. Thus, for a better comprehension of the viral taxonomic distribution, using the same vectors, we filter the data, selecting virus species classified as “examples of ICTV (International Committee on Viral Taxonomy)”^[7]^. A phylogenetic tree was built containing only one virus sample per species, totaling 4,833 viral proteomes (4k).

#### Performance Test

The 31k tree was generated in 2 hours and 17 minutes, and the 4k tree took 13 minutes. rSWeeP had a processing time linear growth curve, as was already expected, of near 500 sequences compared per minute. Thereby, the functions presented by rSWeeP package can give computational power to its users to manage biological big data variables in a practical time.

#### Global dataset analysis

We plotted the two principal components for the two data sets (31k and 4k). We observed that the data presented the same clustering pattern in relation to the type of organization of nucleic acids (Fig. 1). It is evident the separation between the three main clades of single-stranded RNA (ssRNA), single-stranded DNA (ssDNA) and doublestranded DNA (dsDNA) (according to ICTV). We also observed some cases of incompatibility in the genomes, mainly in the ssDNA cluster. These divergences occur by viruses with genotypes that are not inserted in any clade. In addition, there are some cases of taxonomy and annotation errors in the NCBI database, also identified in the study by Calisher and colleagues (2006)^[8]^, and many of these annotations have not been corrected to date. This shows that rSWeeP presents a solution to accurately classify organisms taxonomically.

**Fig. 1.**
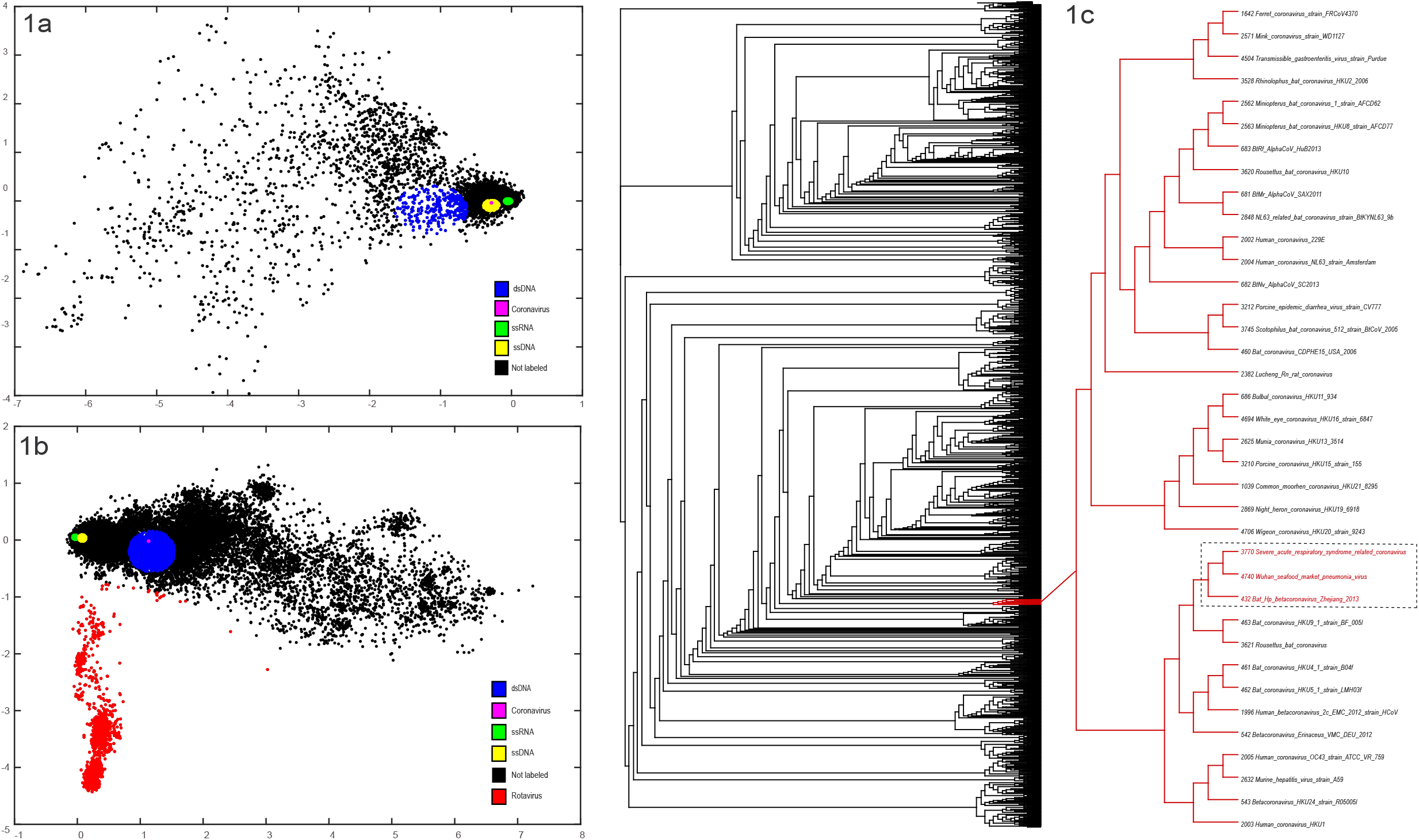
Analysis and representation of virus data sets. 1a. PCA of 4k dataset (4,833 viral proteomes): Single-stranded RNA (green), single-stranded DNA (yellow), double-stranded DNA (blue), Coronavirus (pink), not labeled (black). 1b. Principal component analysis (PCA) of 31k dataset (31,386 viral proteomes): Species with single-stranded RNA (green), single-stranded DNA (yellow), double-stranded DNA (blue), rotavirdae family (red), Coronavirus (pink), not labeled (black)1c. Phylogenetic tree generated using 4k data, with SWeeP default parameters: The enlarged branch containing the coronavirus specimens(red). Featured, the species highly related to SARS-COV1 (square)

#### SARS-CoVs Analysis

According to Gorbalenya and collegues (2020)^[6]^, the criteria adopted to define the coronavirus of the current outbreak, as a “new” coronavirus would not be ideal, since it is the same coronavirus reported by Drostren C. and colleagues (2003)^[9]^. According to the ICTV, the new nomenclatures are SARS-CoV-1 - for the virus isolated in 2003 - and SARS-CoV-2 - for virus isolated in 2019.

For a virus to be considered as a new specimen, it should not be included in known groups, but distant from these groups^[6]^. Our phylogenetic analyzes corroborate this statement. In tree 4k (Figure 1c) we identified the pneumonia strain isolated from the Wuhan seafood market (GCF_009858895.2) - responsible for the current SARS outbreak– is positioned remarkable close to the coronavirus strains related to severe acute respiratory syndrome (GCF_000864885.1) and bat-beta coronavirus (GCF_000926915.1). This reinforces our previous assertation that rSWeeP is effective for analysis and taxonomic classification of viral organisms.

## 4 Concluding Remarks

We show the implementation of SWeeP model in R language, rSWeeP package, is equally accurate. The performance test, phylogenetics and PCA analyzes demonstrate the efficiency of rSWeeP for analysis of viral proteomes. rSWeeP package presents a solution for the taxonomic classification of organisms, freely available for the entire scientific community.

## Acknowledgements

The authors thank Foundation Araucária and the group of Artificial Intelligence applied to Bioinformatics of Federal University of Paraná.

*The authors have declared no conflict of interest.*

